# GDAP1 loss of function inhibits the mitochondrial pyruvate dehydrogenase complex by altering the actin cytoskeleton

**DOI:** 10.1101/2021.03.04.433895

**Authors:** Christina Wolf, Alireza Pouya, Sara Bitar, Annika Pfeiffer, Diones Bueno, Sabine Arndt, Stefan Tenzer, Federica Dal Bello, Caterina Vianello, Sandra Ritz, Jonas Schwirz, Kristina Dobrindt, Michael Peitz, Eva-Maria Hanschmann, Ibrahim Boussaad, Oliver Brüstle, Marta Giacomello, Rejko Krüger, Axel Methner

## Abstract

Charcot-Marie-Tooth (CMT) disease 4A is an autosomal-recessive polyneuropathy caused by mutations of ganglioside-induced differentiation-associated protein 1 (GDAP1), a putative glutathione transferase, which affects mitochondrial shape and alters cellular Ca^2+^ homeostasis. Here, we identify the underlying mechanism. We found that patient-derived motoneurons and *GDAP1* knockdown SH-SY5Y cells display two phenotypes: more tubular mitochondria and a metabolism characterized by glutamine dependence and fewer cytosolic lipid droplets. GDAP1 interacts with the actin-depolymerizing protein Cofilin-1 in a redox-dependent manner, suggesting a role for actin signaling. Consistently, GDAP1 loss causes less F-actin close to mitochondria, which restricts mitochondrial localization of the fission factor dynamin-related protein 1, instigating tubularity. Changes in the actin cytoskeleton also disrupt mitochondria-ER contact sites. This results in lower mitochondrial Ca^2+^ levels and inhibition of the pyruvate dehydrogenase complex, explaining the metabolic changes upon GDAP1 loss of function. Together, these findings reconcile GDAP1-associated phenotypes and implicate disrupted actin signaling in CMT4A pathophysiology.

## Introduction

Charcot-Marie-Tooth (CMT) disease is the most frequently inherited peripheral neuropathy in humans and affects one in 2500 people. Clinically, this group of diseases can be distinguished by mode of inheritance, age of onset, and by electrophysiological characteristics that distinguish demyelinating and axonal forms. Mutations in the gene *GDAP1* (ganglioside-induced differentiation-associated protein 1) cause various forms of CMT: the most frequent recessively inherited demyelinating subtype CMT4A ^1^, the axonal-recessive (AR)-CMT2 ^2^, the intermediate-recessive subtype CMTRIA ^3^, and the dominant subtype CMT2K ^4–6^.

GDAP1 is located in the mitochondrial outer membrane facing the cytosol. It possesses two glutathione transferase (GST)-like domains ^2,7,8^ and a C-terminal hydrophobic anchor which crosses the outer mitochondrial membrane ^9^. Whether GDAP1 is a catalytically active GST remains controversial ^8,10–12^, but it appears clear that GDAP1 can bind GST substrates ^11,12^.

One potential mechanism of GDAP1 action that has been connected to CMT4A disease is mitochondrial dynamics. Over-expression of GDAP1, but not over-expression of GDAP1 bearing recessive disease-causing mutations, results in more fragmented mitochondria, whereas a GDAP1 knockdown (KD) results in mitochondrial elongation ^13^. This GDAP1-mediated mitochondrial fragmentation depends on the activity of dynamin-related protein 1 (DRP1), the major mitochondrial fission factor ^14,15^. The mechanism behind this is unknown. DRP1 is a cytosolic protein which, when recruited to mitochondria, promotes fission by forming ring-like oligomers that constrict and divide mitochondria ^16^. The recruitment of DRP1 to mitochondria depends on the presence of ER tubules ^17^ and filamentous actin (F-actin) ^18^ at the sites of constriction. The ER protein inverted formin 2 (INF2) promotes actin polymerization which occurs before DRP1-driven constriction ^19^. INF2-mediated actin polymerization also increases mitochondria-ER contact sites (MERCS) ^20^. This further connects actin polymerization and GDAP1 function as mitochondrial and ER marker proteins colocalize less frequently in *GDAP1 KD* cells ^21^. Interestingly, INF2 mutations also cause CMT disease (CMTDIE) ^22^, implying that actin polymerization is important for the survival of peripheral nervous system neurons, similar to GDAP1 function. Another protein at the interface of F-actin polymerization and DRP1 recruitment is Cofilin-1. Cofilin-1 binds to monomeric actin and Factin and controls cytoskeletal dynamics mostly by actin depolymerization. Cofilin-1 deletion results in DRP1 accumulation at mitochondria and fragmentation ^23^. In summary, defective actin polymerization affects the same mitochondrial fission pathway upstream of DRP1 and results in a clinical phenotype similar to GDAP1 mutation. If F-actin polymerization and its regulation play a role in CMT4A has not been investigated yet.

Another process that is affected by loss of *GDAP1* is cellular Ca^2+^ homeostasis. Neuronal *GDAP1* KD reduces Ca^2+^ influx from the extracellular space that follows depletion of the ER Ca^2+^ stores, so called store-operated Ca^2+^ entry (SOCE), possibly due to an impaired mitochondrial localization at subplasmalemmal microdomains ^21^. *GDAP1* KD also blunts the Ca^2+^-dependent increase of mitochondrial respiration upon SOCE ^24^. Mitochondrial Ca^2+^ levels increase the activity of several enzymes of the tricarboxylic acid cycle (TCA) like pyruvate dehydrogenase (PDH) ^25^, the key enzyme of the pyruvate dehydrogenase complex (PDC) which catalyzes the conversion of pyruvate to acetyl-CoA and links glycolysis to the TCA. This connection between Ca^2+^ and TCA activity is thought to connect mitochondrial activity to ATP demand. Changes in the activity of the PDC have not been studied yet in CMT4A or in cells with perturbed GDAP1 expression but, interestingly, a pathogenic mutation of pyruvate dehydrogenase kinase isoenzyme 3 (PDK3) that inhibits the PDC also causes CMT disease, CMTX6 ^26^. How GDAP1-mediated changes of the mitochondrial Ca^2+^ homeostasis are connected to its fission activity is still unclear.

In this study, we used motoneurons obtained from CMT4A-patient-derived induced-pluripotent stem cells and neuronal *GDAP1* KD cells to study the pathophysiology of CMT4A. We found that GDAP1 interacts with actin-binding Cofilin-1. Loss of GDAP1 results in a reduction of Factin fibers in mitochondrial proximity, which restricts DRP1 access to mitochondrial constriction sites and disrupts mitochondria-ER contact sites. This reduces mitochondrial Ca^2+^ levels and inhibits the PDC resulting in a rewired cellular metabolism characterized by glutamine dependence and increased consumption of fatty acids. Together, these findings implicate disrupted F-actin signaling in CMT4A pathophysiology.

## Results

### More tubular mitochondria and an increased mitochondrial membrane potential in CMT4A patient-derived neuronal cells and GDAP1 knockdown cells

To establish a model for CMT4A, we compared *GDAP1* KD SH-SY5Y cells ^21,27^ with neuronal cells derived from CMT4A patients. Patient CMT#1 is a 25-year-old ambulant male with a compound heterozygosity (L239F/R273G) of mutations in the C-terminal GST domain of GDAP1 and patient CMT#2 is a 40-year-old wheelchair-bound male with a homozygous mutation of the intron 4 splice donor site (c.579 + 1G>A). This mutation causes skipping of exon 4 leading to a frameshift and a truncated protein lacking the C-terminal GST and the transmembrane domain of GDAP1 (Fig. 1a).

**Figure 1.**
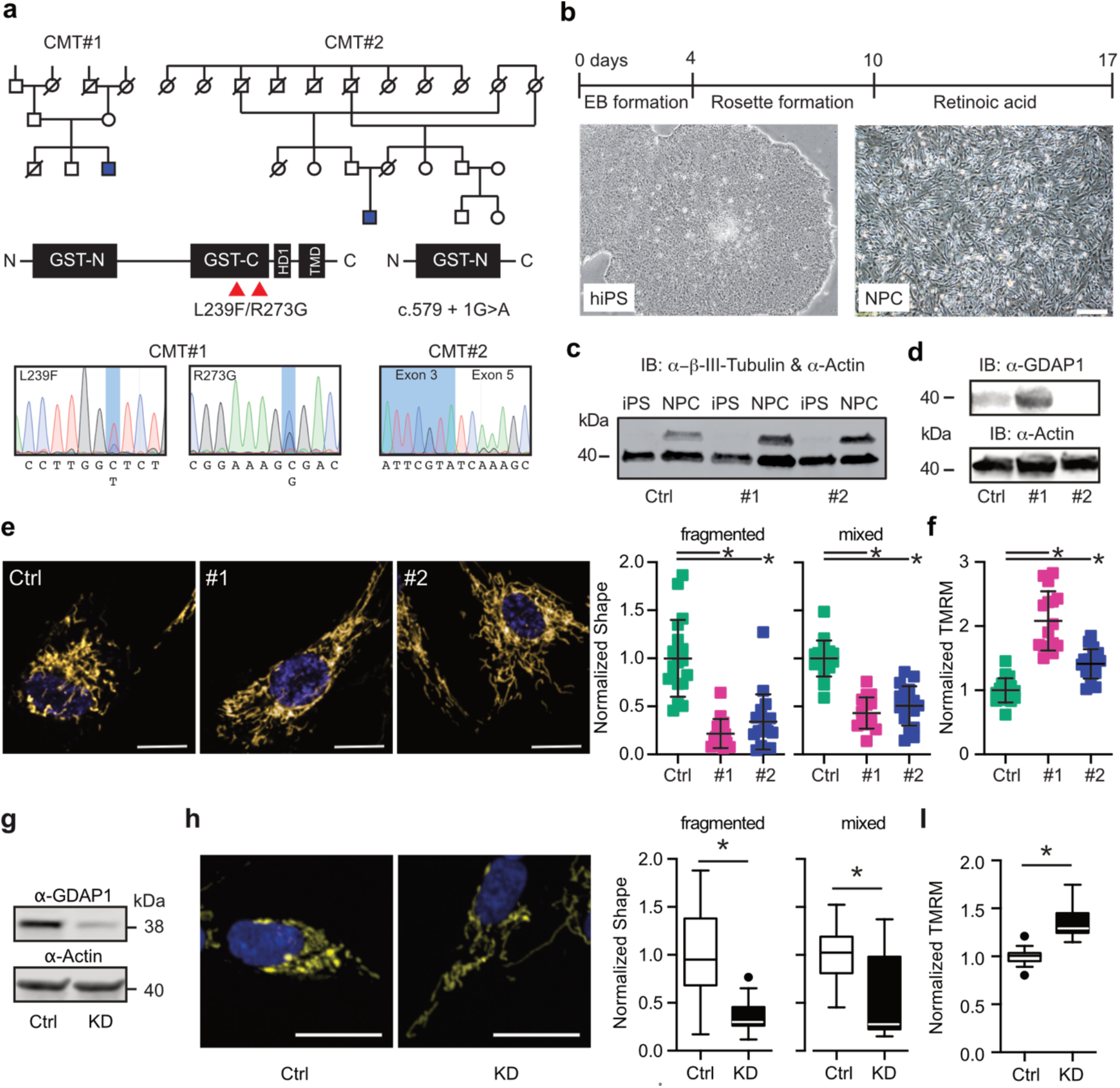
More tubular mitochondria and an increased mitochondrial membrane potential in CMT4A patient-derived neuronal cells and GDAP1 knockdown cells. **(a)** Pedigree and GDAP1 DNA sequences from two patients suffering from autosomal-recessive CMT4A disease. **(b)** Differentiation protocol to obtain neuronal precursor cells from induced pluripotent stem cells (iPSCs), scale bar: 200 μm. **(c)** Immunoblot demonstrating expression of the neuronal marker β-tubulin III in the NPCs but not in iPSCs. **(d)** Control and CMT#1 but not CMT#2 neuronal cells express GDAP1 shown by immunoblotting. Size is indicated, Actin served as loading control. **(e/h)** Representative images of automated high-content confocal microscopy analysis of mitochondrial shape (MitoTracker, scale bar 20 μm) and **(f/i)** mitochondrial membrane potential (TMRM) demonstrating elongated mitochondria and increased mitochondrial membrane potential in CMT4A NPCs and GDAP1 knockdown (KD) cells. **(g)** GDAP1 immunoblot demonstrating successful knockdown. Size is indicated, Actin served as loading control. Data in **e** and **f** are from 3 independent experiments with 4-8 replicates per experiment with a range of 74 to 1908 cells per well. Data in **h** and **i** are from 4 independent experiments with 4-8 replicates with a range of 343 to 4977 cells per well. Statistical variation is shown as scatter plot (**e, f**) or Tukey boxplot (**h, i**) and significance calculated using the non-parametric Kruskal-Wallis (**e, f**) or Mann-Whitney (**h, i**) tests, *p<0.05.

We generated induced-pluripotent stem cells from fibroblast cell lines ^28^ from these patients using non-integrative expression of the Yamanaka factors and differentiated them to neuronal precursor cells (NPCs) (Fig. 1b) that express the neuronal marker protein β-III tubulin (Fig. 1c). In contrast to fibroblasts ^28^, control and CMT#1 NPCs express GDAP1 detectable by immunoblotting (Fig. 1d). NPCs from CMT#2 lacked GDAP1 expression. Because the polyclonal antiserum targets an antigen which should still be present in the patient, this is probably due to nonsense-mediated mRNA decay or degradation of the truncated protein.

We quantified mitochondrial shape and membrane potential (Δψm) using automated high-content confocal microscopy analysis of cells simultaneously stained with the fluorescent dyes mitotracker and tetramethylrhodamine methyl ester (TMRM). Patient-derived neuronal cells contained significantly more tubular mitochondria (Fig. 1e) with a more negative mitochondrial membrane potential (Fig. 1f). Using the same methodology with *GDAP1* KD cells (Fig. 1g), we found similar changes; more elongated mitochondria (Fig. 1h) and a significantly more negative membrane potential (Fig. 1i). The consistency between the phenotypes of patient-derived cells and *GDAP1* KD cells suggests that *GDAP1* KD cells can be used as a model to study CMT4A disease.

### GDAP1 knockdown uncouples mitochondrial respiration from ATP generation

Mitochondria produce ATP by consuming oxygen and the oxidizing agent NAD+ in a process called oxidative phosphorylation. NAD+ is provided by the TCA. To assess the effect of *GDAP1* KD on this all-important process, we measured mitochondrial oxygen consumption using high-resolution respirometry and found a higher mitochondrial routine respiration in *GDAP1* KD cells (Fig. 2a). The maximal capacity of the electron transfer system (ETS), determined by titrating in the uncoupler FCCP, was however similar in both cell lines (Fig. 2a). Normalization of the respiratory states to the maximum ETS capacity revealed that the *GDAP1* KD cells use a higher fraction of their maximal capacity for routine, leak and phosphorylating respiration (Fig. 2a’). However, quantification of ATP content using the ratiometric reporters BTeam targeted either to the cytosol or to the mitochondrial matrix ^29^ demonstrated a reduced mitochondrial ATP generation in *GDAP1* KD cells despite the increased Δψm and routine respiration (Fig. 2b). Taken together, these data suggest that *GDAP1* KD results in the uncoupling of mitochondrial respiration and ATP generation via oxidative phosphorylation.

**Figure 2.**
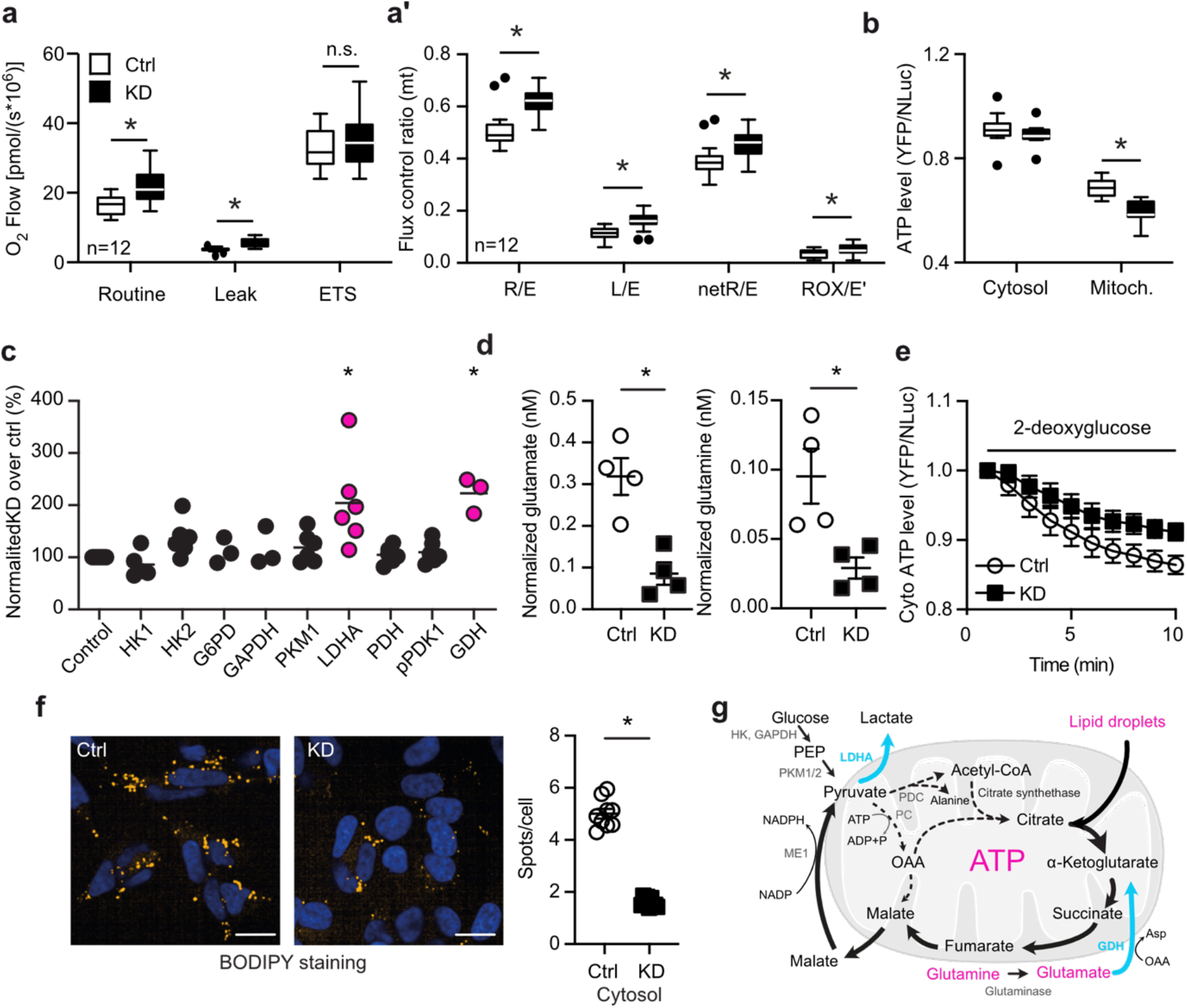
GDAP1 knockdown uncouples mitochondrial respiration from ATP generation and shifts metabolism towards glutaminolysis. **(a)** Mitochondrial oxygen consumption of intact cells in regular growth medium measured by high-resolution respirometry and corrected for ROX. ETS, electron transfer system; L, leak respiration; R, routine respiration; E, ETS capacity; ROX, residual oxygen consumption. **(a’)** Calculation of flux control ratios. **(b)** Comparison of cytosolic and mitochondrial ATP content using the genetically encoded ATP sensor BTeam. Calculation of BTeam YFP/NLuc emission ratios under basal conditions revealed reduced mitochondrial ATP levels. **(c)** Immunoblot quantification of proteins involved in glycolysis in Ctrl and KD cells shows an increase in LDHA and GDH levels. HK, Hexokinase; G6PD, glucose-6-phosphate dehydrogenase; GAPDH, Glyceraldehyde 3-phosphate dehydrogenase; PKM1, pyruvate kinase M1; LDHA, lactate dehydrogenase A; PDH, pyruvate dehydrogenase; pPDK1, phospho-pyruvate dehydrogenase kinase; GDH, glutamate dehydrogenase. Actin expression served to normalize the data. **(d)** Diminished glutamate and glutamine levels determined by a luminescence-based assay. **(e)** Comparison of cytosolic ATP content using the genetically-encoded ATP sensor BTeam. BTeam YFP/NLuc emission ratios after treatment with 25 μM 2-deoxyglucose (2-DG) reveals increased non-mitochondrial ATP generation capacity in KD cells. **(f)** Automated high-content confocal microscopy analysis of BODIPY-stained fatty acids demonstrating less lipid droplets in KD cells identified by Höchst staining of nuclei. **(g)** Schematic illustration of metabolic changes observed in *GDAP1* KD cells. Upregulation is shown in blue and bold lines; downregulation in magenta and dashed lines. ME, malic enzyme 1; PDC, pyruvate dehydrogenase complex; PKM1/2, pyruvate kinase M1/2; OAA, oxaloacetate. Data in **a** and **a’** are from 12 independent experiments performed in duplicate. Data in **b** are from 4 independent experiments performed in triplicates. Data in **d** are from 4 independent experiments performed in triplicates. Data in **e** are from 6 independent experiments performed in triplicates. Data in **f** were from >10000 cells in total and were analyzed in 3 independent experiments, performed in triplicates each. Statistical variation is shown as Tukey’s boxplot, XY graph or scatter plot with the indication of the mean ± SEM. Significance was calculated using the non-parametric Kruskal-Wallis test, *p<0.05.

### GDAP1 knockdown shifts cellular metabolism towards glutaminolysis

To identify metabolic pathways in *GDAP1* KD cells that compensate for the diminished capacity for oxidative phosphorylation, we quantified the expression levels of various metabolic enzymes in control and KD cells by immunoblotting (Supplementary Fig. 1). This revealed a significant upregulation of lactate dehydrogenase (LDHA) and glutamate dehydrogenase (GDH) in *GDAP1* KD cells (Fig. 2c). LDHA catalyzes the interconversion of pyruvate to lactate. GDH, in contrast, converts glutamate to α-ketoglutarate, the precursor of succinyl-CoA in the TCA. In line with the increased expression of GDH, KD cells consume significantly more glutamate, the substrate of GDH, and more glutamine which can be converted to glutamate by glutaminase (Fig. 2d). We also observed an attenuated decline in cytosolic ATP content in KD cells after treatment with the glucose antimetabolite 2-deoxyglucose (Fig. 2e) in line with a reduced use of glucose as the major fuel for the TCA cycle. The TCA metabolite upstream of α-ketoglutarate is citrate, which is produced by the transfer of acetyl-CoA to oxaloacetate. Acetyl-CoA is the product of the PDC in the mitochondrial matrix or of mitochondrial β-oxidation. We therefore quantified the amount of lipid droplets by BODIPY C12 staining and found decreased cytosolic levels in *GDAP1* KD cells (Fig. 2f) suggesting a high lipid catabolism that serves to replenish fatty acids and consequently acetyl-CoA levels in the TCA cycle. The increased demand of glutamine and fatty acids in *GDAP1* KD probably serve as compensatory mechanisms, clearly pointing towards a dysfunction of the TCA cycle in *GDAP1* KD cells at the level of the PDC (summarized in Fig. 2g).

### Hyperphosphorylated pyruvate dehydrogenase and reduced mitochondrial Ca^2+^ levels in GDAP1 KD cells

Increased mitochondrial Ca^2+^ levels activate the PDC by stimulating PDH phosphatase ^30^. Ca^2+^ also activates isocitrate dehydrogenase which is upstream of α-ketoglutarate, the product of GDH, and α-ketoglutarate dehydrogenase ^31^. Based on the reported attenuated mitochondrial respiration upon SOCE in *GDAP1* KD cells ^24^, we suspected altered PDH phosphorylation levels driven by changes in mitochondrial Ca^2+^ content as the reason for PDC inhibition (see scheme in Fig. 3a). To test this idea, we probed phosphorylation of serine 293 of the E1 PDH subunit, which has been directly linked to the PDC activity ^32^. Immunoblotting showed a significantly increased phosphorylation of PDH E1 serine 293 in *GDAP1* KD cells as compared to total E1 PDH (Fig. 3b). This is in line with a decreased activity of the PDC. We then measured mitochondrial Ca^2+^ levels by imaging live cells stained with Rhod2-AM or expressing the genetically-encoded mitochondrial Ca^2+^ sensor mito-CEPIA ^33^ normalized to mito-FarRed. The Ca^2+^ sensor GEM-CEPIA1er targeted to the ER served as a control. Both methods revealed a reduction in steady-state mitochondrial Ca^2+^ levels in *GDAP1* KD cells, whereas ER calcium levels were not affected (Fig. 3c). We conclude that the PDC malfunction in *GDAP1* KD cells is likely caused by a reduction in mitochondrial Ca^2+^ levels resulting in an increased phosphorylation of PDH.

**Figure 3.**
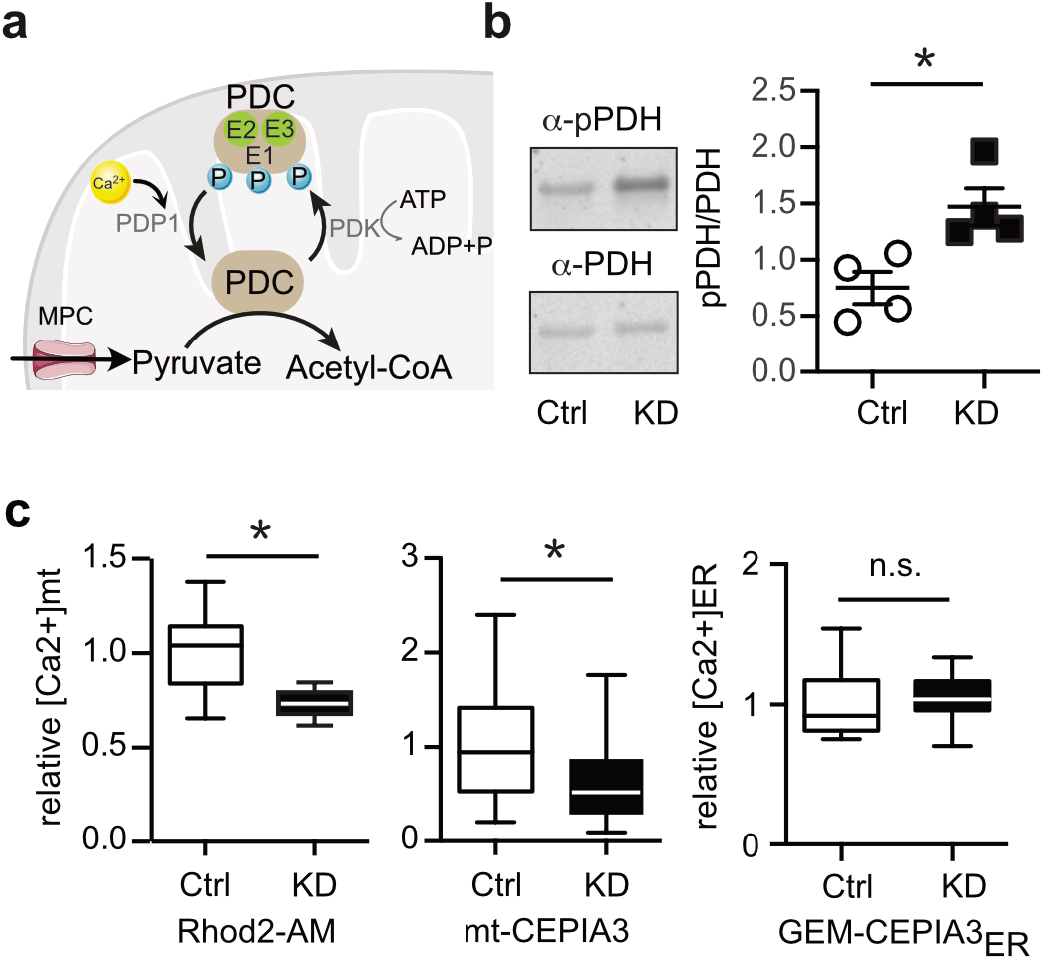
Hyperphosphorylated pyruvate dehydrogenase and reduced mitochondrial Ca^2+^ levels in GDAP1 KD cells. **(a)** Scheme showing the regulation of the pyruvate dehydrogenase complex (PDC). The PDC E1 subunit can be phosphorylated by the catalytic activity of the PDH kinase (PDK). The PDH phosphatase subunit 1 (PDP1) in turn dephosphorylates the serine residues upon activation by Ca^2+^. MPC, mitochondrial pyruvate carrier; PDK1, pyruvate dehydrogenase kinase 1. **(b)** Immunoblots from whole cell lysates for quantification of PDH E1 phosphorylation (serine 293), normalized to total PDH E1 levels revealed increased phosphorylation of PDH in *GDAP1* KD cells. **(c)** Mitochondrial Ca^2+^ measured with the fluorescent dye Rhod2-AM or mito-CEPIA normalized to mito-FarRed indicated reduced mt[Ca^2+^] levels. ER[Ca^2+^] levels measured with the genetically encoded Ca^2+^ sensor GEM-CEPIA3_ER_ did not show any variation between the cell lines. Data in **c** were from 3 independent experiments of n=35 cells (KD), n=44 cells (Ctrl) for the Rhod2-AM analysis and n=26-28 (KD) respectively n=15-34 cells (Ctrl) for mito-CEPIA3 analysis. Data for GEM-CEPIA3_ER_ were from 2 independent experiments, KD, n=27-30 and Ctrl 29-45 cells. Statistical variation is shown as Tukey boxplots or mean ± SEM and significance calculated using the non-parametric Mann-Whitney, *p<0.05.

### Patient-derived cells are similarly characterized by an anaplerotic state and reduced mitochondrial Ca^2+^ levels

We next tested whether the findings of reduced mitochondrial Ca^2+^ levels and increased glutaminolysis also apply to patient-derived cells. We set out to specifically test this in motoneurons, a cell type affected by CMT4A. Motoneurons were differentiated from NPCs following established protocols (Fig. 4a) and motoneuronal identity was confirmed by immunostaining against the dendrite marker MAP2 and the motoneuronal neurofilament H marker antibody Smi32 (Fig. 4b). These motoneurons express GDAP1 similar to NPCs with cells from patient CMT#2 lacking GDAP1 expression (Fig. 4c). High-content microscopy of MitoTracker-stained cells revealed an increased area occupied by mitochondria (Fig. 4d). Immunoblotting reproduced the increased expression of GDH seen upon *GDAP1* KD (Fig. 4e, Fig. 2c). Motoneurons differentiated from patient-derived cells also increased the consumption of the GDH-substrate glutamate and its precursor glutamine (Fig. 4f, Fig. 2d). Furthermore, motoneurons showed a decrease in lipid droplets normalized to the nuclear area (Fig. 4g), indicative of increased fatty acid consumption. Because motoneuronal cells showed a low transfection efficiency which made it difficult to identify single neurons in the cultures, we reverted to the patient-derived NPCs to assess the mitochondrial Ca^2+^ phenotype. We measured lower resting Ca^2+^ levels with Rhod2-AM and mito-CEPIA (Fig. 4h) in NPCs, similar to KD cells. We conclude that motoneurons derived from patients with GDAP1 mutation demonstrate the same metabolic changes and alterations of Ca^2+^ levels identified in *GDAP1* KD cells, strengthening the evidence that our findings reveal the mechanism underlying GDAP1-mediated CMT4A.

**Figure 4.**
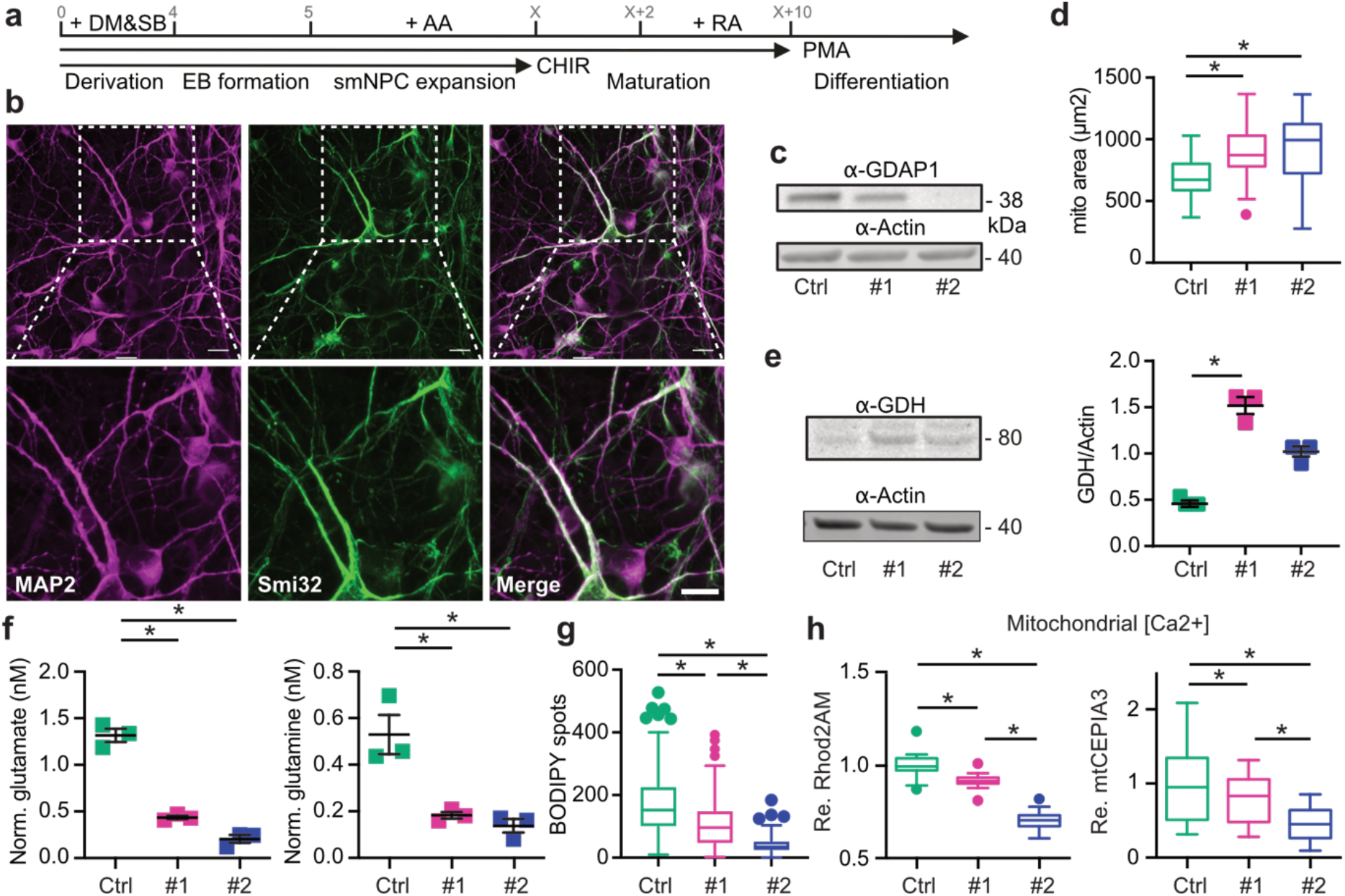
Patient-derived cells are similarly characterized by an anaplerotic state and reduced mitochondrial Ca^2+^ levels. **(a)** Differentiation protocol to obtain motoneurons (MN) from induced pluripotent stem cells (iPSCs) and **(b)** confirmatory immunocytochemistry by staining against dendrite marker MAP2 and motoneuronal neurofilament Smi32. **(c)** Immunoblot of total MN cell lysates for validation of GDAP1 expression, actin served as loading control, size is indicated. **(d)** Automated high-content microscopy analysis of MitoTracker-stained mitochondria revealed an increased total mitochondrial mass per well. **(e)** Immunoblot of total MN cell lysates against glutamate dehydrogenase (GDH), with actin as loading control, and **(f)** reduced glutamate and glutamine levels in CMT4A patient-derived MN determined by a luminescence-based assay indicating increased glutamine consumption compared to Ctrl. **(g)** Automated high-content microscopy analysis of BODIPY-positive spots normalized to the nuclei area in MN revealed a reduced number of lipid droplets in CMT4A patient-derived cells. **(h)** Reduced relative mt[Ca^2+^] levels of NPCs determined by confocal microscopy with Rhod2-AM and mito-CEPIA3. Data in **d-g** were from 3 independent differentiations and data in **d,g** were obtained from n>40,000 cells in quadruplicate, experiments in f were performed in duplicate. Data in **h** were from 20-22 Rhod2-AM-stained cells and 22-33 cells mitoCEPIA3 transfected from three independent experiments. Statistical variation is shown as Tukey boxplots or scatter plots with mean ± SEM indicated. Significance was calculated using the non-parametric Kruskal Wallis test, *p<0.05.

### GDAP1 knockdown reduces the number of contact sites between mitochondria and the endoplasmic reticulum

We suspected that changes in MERCS might underlie the changes in mitochondrial Ca^2+^ levels especially as a reduced colocalization of ER and mitochondrial markers as a hint to changes in MERCS was already shown previously ^21^. MERCS are hot spots of interactions between the ER and mitochondria and important signaling hubs (reviewed in ^34,35^). Using transmission electron microscopy, we indeed found i) a decrease in MERCS in *GDAP1* KD cells defined as the percentage of mitochondrial perimeter covered by the ER, ii) a decreased number of MERCS per mitochondrion, iii) a decreased length of MERCS and iv) an increased width between the ER and mitochondria (Fig. 5a). This leads to a complete right shift of MERCS width in *GDAP1* KD cells (Fig. 5a’). Comparing the amount of ER protein present in mitochondrial cell fractions using immunoblotting of the membrane ER protein Sec62 and the mitochondrial protein Cox4 confirmed the reduction in contact sites (Fig. 5b). As an additional readout to quantify the distance between mitochondria and the ER, we used a well-established assay based on Förster resonance emission transfer (FRET) between two fluorescent proteins targeted to the surface of mitochondria and the ER, both facing the cytosol ^36,37^. As FRET only occurs when the two fluorophores are close enough, the lower FRET ratio (Fig. 5c) confirmed the data obtained by the morphometric analysis with an increased width between the organelles in *GDAP1* KD cells. Such an increased distance should result in decreased Ca^2+^ transfer from the ER to the mitochondrial matrix. We studied this by triggering ER Ca^2+^ release with the cholinergic agent carbachol, which activates inositol trisphosphate receptors (IP3Rs) leading to Ca^2+^ release into the cytosol, generating Ca^2+^ microdomains at MERCS, which sustain rapid Ca^2+^ uptake by mitochondria ^36,38^. Surprisingly, we did not observe a difference in mitochondrial Ca^2+^ uptake measured by mitochondrially-targeted aequorin upon treatment with carbachol (Fig. 5d). The direct reducing effect of carbachol on PDH E1 phosphorylation was also similar in both cell lines while the baseline levels were significantly increased in *GDAP1* KD cells (Fig. 5e). Apparently, upregulated MCU levels (Fig. 5f) and hyperpolarization (see Fig. 1I) can compensate for the increased distance between the organelles under conditions of high Ca^2+^ influx. Taken together our findings support altered MERCS with an increased distance between the ER and mitochondria in *GDAP1* KD cells.

**Figure 5.**
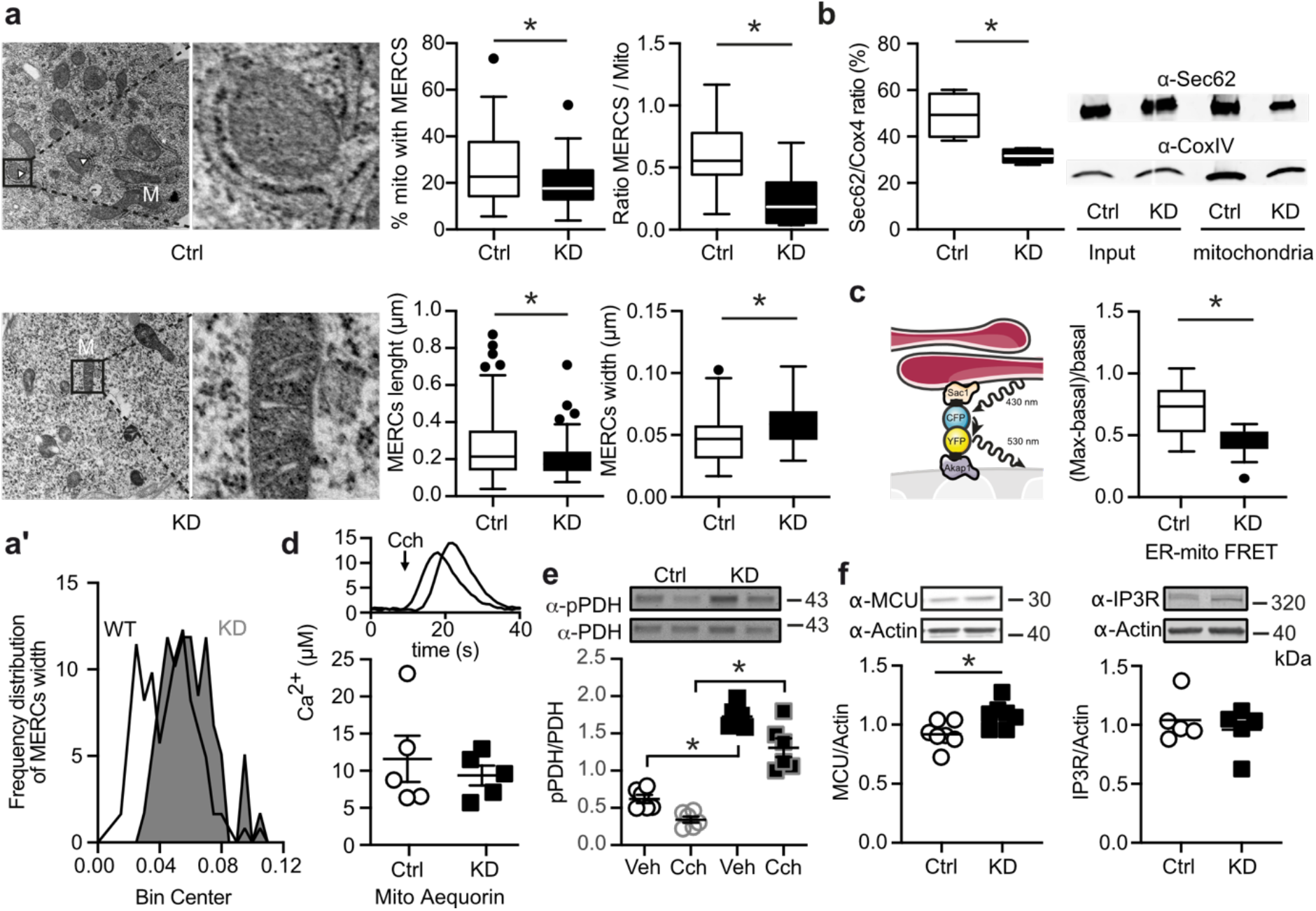
GDAP1 knockdown reduces the number of contact sites between mitochondria and the endoplasmic reticulum. **(a)** Transmission electron microscopy of knockdown (KD) and control (Ctrl) cells. The indicated parameters were quantified using ImageJ by a blinded investigator. Scale bar in A refers to 1000 nm; scale bar in respective magnifications refers to 500 nm. **(a’)** Histogram showing the distribution of MERCS’ widths, and an increased distance in KD cells between the ER and mitochondria. **(b)** Quantification of ER membrane (Sec62) and mitochondrial (Cox4) proteins in mitochondrial fractions. **(c)** Scheme of the FRET-FEMP sensor comprising ER CFP-Sac1 and mitochondrial YFP-Akap1. Proximity of both organelles leads to high intensity of YFP-FRET-emission (410-430/520-560 ex/em). 100 nM rapamycine was added to achieve the closest possible distance, and the measurement of FRET ratio was calculated as (FRET_max_-FRET_basal_)/FRET_basal_. **(d)** Representative curve for the aequorin Ca^2+^ measurement and quantification of mitochondrial Ca^2+^ levels after addition of 200 μM carbachol (CCH), which releases Ca^2+^ from the ER by activation of a G-protein-coupled receptor. **(e)** Immunoblot of Ctrl and KD cells lysed 10 minutes after 200 μM CCH addition to quantify the PDH phosphorylation (pPDH/(PDH_total_) after Ca^2+^ release into the MERCS and mitochondria. **(f)** Immunoblot of total cell lysates of Ctrl and KD cells showing increased expression levels of MCU but not pan-IP3R, actin served as loading control, size is indicated. Data in **a** were obtained from 11 (Ctrl) and 9 (KD) cells, in **b** from 4 independent experiments, and data in **c** were from 4 independent experiments in triplicate or quintuplicate with a range of 162-818 cells per experiment. Data in **d** from 5 independent experiments performed in triplicates. Statistical variation is shown as Tukey boxplots or scatter plots with the indication of mean ± SEM and significance was calculated using the non-parametric Mann-Whitney test, *p<0.05.

### GDAP1 interacts with the actin-depolymerizing protein Cofilin-1 in a redox-dependent manner

To clarify how *GDAP1* KD affects MERCS and mitochondrial Ca^2+^ levels, we next set out to identify interacting proteins by label-free quantitative mass spectrometry ^39,40^. We transduced primary cortical cultures from mice expressing the biotin ligase BirA ^41^ with adeno-associated viruses expressing GDAP1 tagged with an Avi-tag, a specific substrate for BirA ^42,43^. Only Avi-HA-tagged GDAP1 but not the HA-tagged control was biotinylated (Fig. 6a). By treating the mitochondrial preparations with GSH or GSSG as described ^44^, we further aimed to identify proteins whose interaction is affected by their specific redox state. This was driven by the fact that GDAP1 can bind potential GST substrates ^11,12^ and the assumption that such an interaction might depend on the local redox environment. A total of 163 proteins were specifically pulled down from Avi-HA GDAP1-transduced cultures (Fig. 6b). Out of these, 60 proteins showed a different abundance in preparations treated with GSH and GSSG. Interestingly, many of these proteins can be linked to the cytoskeleton like tubulins (Fig. 6c, green), crucial components of the cytoskeleton that can serve as a scaffold for mitochondrial transport ^45,46^. Interestingly, an interaction between GDAP1 and β-tubulin TUBB has been reported in a yeast-two-hybrid experiment ^47^. The protein Cofilin-1 specifically caught our attention because it affects mitochondrial shape by inducing depolymerization of actin filaments in mitochondrial proximity which restricts access of DRP1 to mitochondria ^23^. Its antagonist is the ER-anchored INF2, which also causes CMT disease when mutated ^22^.

**Figure 6.**
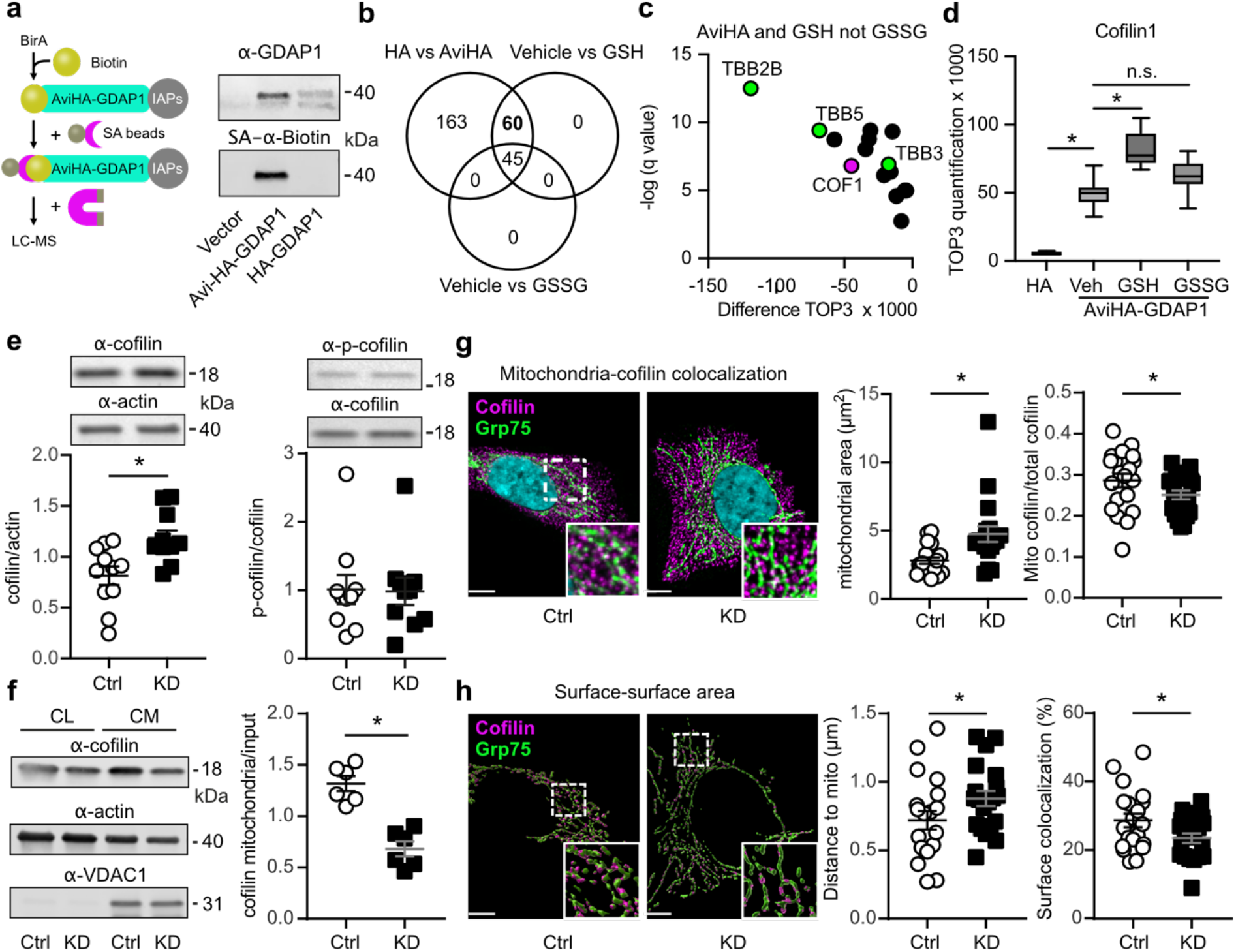
GDAP1 interacts with the actin-depolymerizing protein Cofilin-1 in a redox-dependent manner and reduces Cofilin-1 abundance at mitochondria. **(a)** Schematic illustration showing the biotinylation of the AviTag fused to GDAP1 protein and subsequent pulldown and enrichment of biotinylated GDAP1 protein via streptavidin-labeled magnetic beads. Immunoblot of cell lysates prior to pulldown of biotinylated GDAP1 stained against GDAP1 and Biotin with an infrared-labelled streptavidin (SA) dye shows an overlap of biotinylation and GDAP1 protein. BirA pulldown was performed with neurons isolated from E16 embryos. **(b-d)** Results from label-free quantitative proteomics. TOP3 quantification of Cofilin-1 abundance and increased Cofilin-1 abundance upon GSH, but not GSSG addition indicated an interaction with GDAP1 in a redox-dependent manner. **(e)** Immunoblot of whole cell lysates and quantification of Cofilin-1 and p-Cofilin-1 (S3) protein levels. Actin served as loading control. **(f)** Analysis of Cofilin-1 abundance at the mitochondria after fractionation of crude mitochondria (CM) normalized to Cofilin-1 levels in whole cell lysates (CL). **(g)** Immunostaining of Ctrl and GDAP1 KD cells for Cofilin-1 (magenta) and GRP75 (green) as mitochondrial protein. Nuclei were counterstained with DAPI. Quantification of the mitochondrial area confirmed the increased mitochondrial size in KD cells and mitochondrially-located Cofilin-1 was significantly reduced. **(h)** Conversion of the structures into surfaces using Imaris software allowed evaluation of the contact area (magenta). Analysis of the distance between Cofilin-1 spots to the closest mitochondrion also revealed an increased distance in GDAP1 KD cells. Imaris surface contact XTension was used to determine the proportion of the mitochondrial surface area in contact with Cofilin-1, which was significantly reduced. **(g,h)** Scale bar equals 5 μm; n=20 cells per cell line. Data from **b-e** were from three independent experiments, which were measured in quadruplicates each. Statistical significance was determined by t-test with Bonferroni correction. Data in **g-h** are from n=20 cells per cell line performed in triplicates. Statistical variation is shown as Tukey boxplots or scatter plots with the indication of mean ± SEM and significance calculated using the non-parametric Mann-Whitney test, *p<0.05.

### GDAP1 knockdown reduces Cofilin-1 abundance at mitochondria

We further explored the identified interaction between GDAP1 and Cofilin-1, and hypothesized that GDAP1 alters actin abundance, polymerization or both at mitochondrial sites by interacting with Cofilin-1. The actin-binding ability of Cofilin-1 is inhibited by phosphorylation of serine 3 which is controlled by the presence of intramolecular disulfide bridges ^48,49^ or protein glutathionylation ^50^ and could thus be a target of GDAP1’s GST activity. Alternatively, GDAP1 could just recruit Cofilin-1 to the mitochondrial surface in a redox-dependent manner. We studied Cofilin-1 abundance and intracellular localization and phosphorylation in *GDAP1* KD and control cells. *GDAP1* KD increased the total abundance of Cofilin-1 but did not affect phosphorylation (Fig. 6e). However, when we compared the protein levels in mitochondrial fractions, we observed the opposite, a significant reduction in *GDAP1* KD cells (Fig. 6f), which was also evident when we quantified the intracellular amount and localization of Cofilin-1 close to mitochondria using confocal image analysis. Mitochondrial mass was increased, whereas Cofilin-1 at the mitochondria was reduced in *GDAP1* KD cells (Fig. 6g). In addition, the distance of Cofilin-1 spots from mitochondria was significantly increased and the co-localization of the surfaces of Cofilin-1 and Grp75-expressing structures was reduced (Fig. 6h). These results imply that GDAP1 controls the presence of Cofilin-1 in proximity to the mitochondrial surface.

### GDAP1 KD reduces F-actin fibers at mitochondrial surfaces and restricts access of DRP1 to the mitochondria

Because Cofilin-1 is an actin-binding protein, we suspected less actin in the proximity of mitochondria in *GDAP1* KD cells. We quantitated F-actin levels and its colocalization with mitochondria in living cells by transiently transfecting control and *GDAP1* KD cells with GFP-tagged F-tractin and labeling of mitochondria with mitotracker red. In contrast to other methods like phalloidin staining, F-tractin does not perturb actin rearrangement ^51^ and was previously used to study the effects of INF2 perturbation on mitochondrial dynamics ^20,52^. Our data revealed no changes in the distance between mitochondria and F-actin (Fig. 7a) but a significant reduction of surface colocalization between F-actin and mitochondria in *GDAP1* KD cells (Fig. 7b) in line with our hypothesis. As expected from previous work ^19^, this severely reduced DRP1 localization in mitochondrial fractions (Fig. 7c) explaining the increased tubular shape of mitochondria in GDAP1 loss-of-function cells. These findings imply that in the absence of GDAP1, F-actin fibers are less present at the mitochondrial surface which restricts the access of ER tubules and DRP1 to sites of mitochondrial constriction resulting in a dysfunction of mitochondrial dynamics (Fig. 7d).

**Figure 7.**
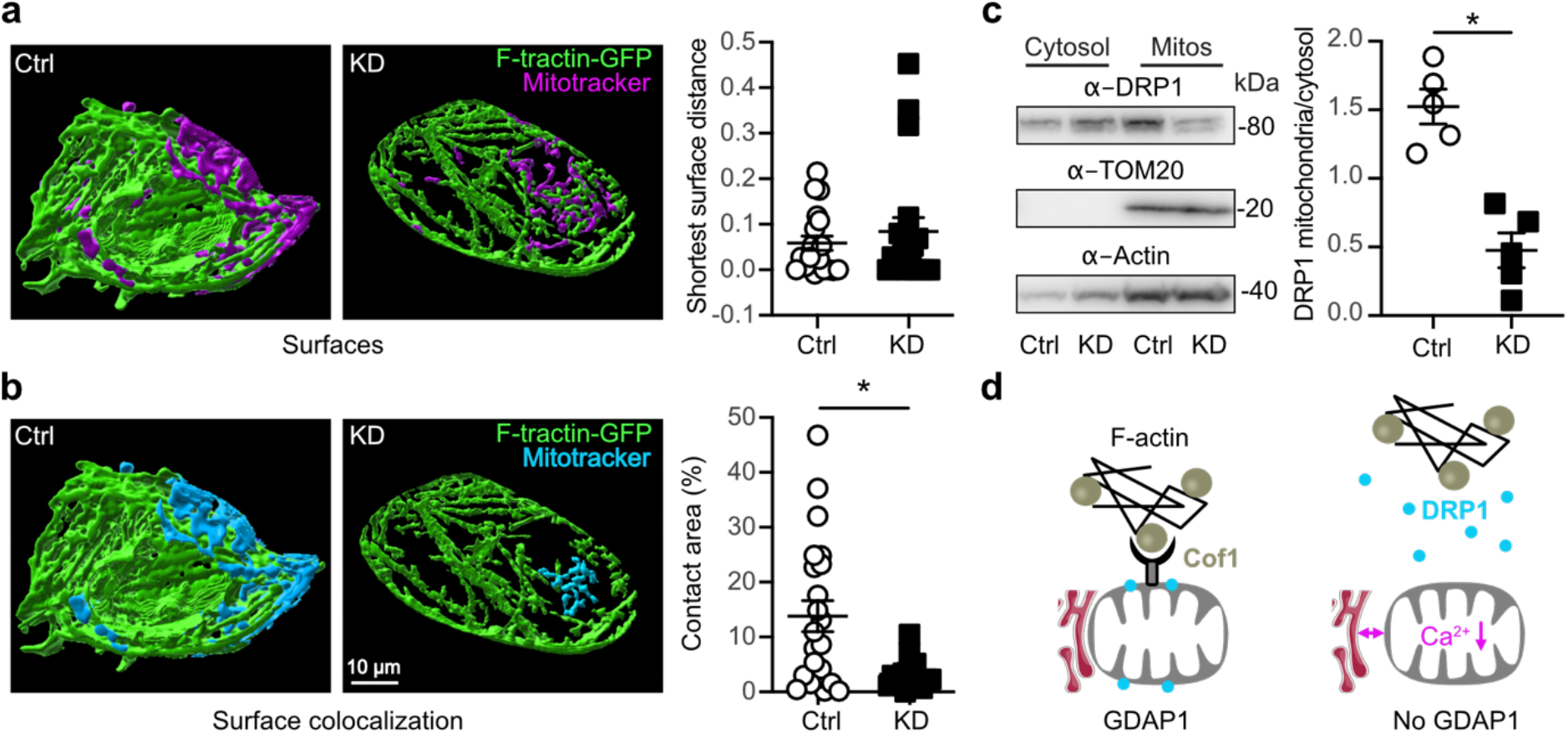
GDAP1 KD reduces F-actin fibers at mitochondrial surfaces and restricts access of DRP1 to the mitochondria. **(a/b)** Live cell imaging of GFP-F-tractin transfected cells. Mitochondria were stained with MitoTracker and images analyzed for the **(a)** distance and **(b)** colocalization of actin filaments and mitochondria, which demonstrated a reduced surface colocalization in *GDAP1* KD cells. **(c)** Immunoblot of DRP1 in cytosolic and mitochondrial fractions demonstrating a reduced DRP1 abundance in mitochondrial fractions. TOM20 and actin served as loading controls, size is indicated. **(d)** Scheme depicting our findings. In the absence of GDAP1 mitochondrial F-actin is reduced resulting in an increased distance between ER and mitochondria, reduced mitochondrial Ca^2+^ levels, and a reduced presence of DRP1 at mitochondrial constriction sites. Data from **a** and **b** are from n=19 cells per cell line from three independent experiments performed in triplicates. Statistical variation is shown as scatter plots with the indication of mean ± SEM and significance calculated using the non-parametric Mann-Whitney test, *p<0.05

## Discussion

In this work, we used neuronal *GDAP1* KD in human SH-SY5Y neuroblastoma cells and patient-derived cells to study the pathophysiology of CMT4A. We found that GDAP1 interacts with the actin-interacting protein Cofilin-1 and that loss of GDAP1 results in a reduction of Factin fibers in mitochondrial proximity. This limits the access of ER tubules to mitochondria which connects two processes: it impedes DRP1 recruitment to mitochondrial constriction sites resulting in more tubular mitochondria and it disrupts mitochondria-ER contact sites causing reduced mitochondrial Ca^2+^ levels. The reduced mitochondrial Ca^2+^ levels inhibit the PDC and result in a rewired cellular metabolism characterized by glutamine and fatty acids dependence to compensate for an impaired TCA cycle. We therefore conclude that the reduction in F-actin presence at mitochondria caused by GDAP1 loss of function represents the probable cause of autosomal-recessive CMT4A.

We found that GDAP1 interacts with Cofilin-1 in a redox-dependent manner and restricts its presence at the mitochondrial surface. Theoretically, GDAP1 could also affect Cofilin-1 function through its still unresolved potential GST-like enzymatic activity ^8,10–12^ because Cofilin-1 contains four potential GST target cysteine residues at the positions 39, 80, 139 and 147. Cysteines 39 and 80 are buried inside the protein while cysteines 139 and 147 are located on the surface of the protein ^53^. Redox-mediated modifications of these cysteine residues clearly affect the function of Cofilin-1. Treatment with hydrogen peroxide leads to the formation of an intramolecular disulfide bond resulting in a conformational change that prevents phosphorylation and thereby actin polymerization ^48^. Moreover, intermolecular disulfides and oligomeric forms of Cofilin-1 have been described. Monomeric Cofilin-1 possesses severing activity, whereas the dimeric and oligomeric forms have actin-bundling activity ^54^. Interestingly, only monomeric Cofilin-1 is phosphorylated ^55^ which represents the most-studied post-translational modification of Cofilin-1 to date. Cofilin-1 can be post-translationally modified by phosphorylation of serine 3. Increased phosphorylation inhibits the actin-binding ability of Cofilin-1 ^56^; dephosphorylated Cofilin-1 preferentially localizes to mitochondria, whereas a mutation mimicking the phosphorylated protein prevents translocation from the cytosol to mitochondria ^57^. We found no changes in Cofilin-1 phosphorylation in whole cell lysates and no phosphorylated Cofilin-1 in mitochondrial fractions. Upon oxidation of all four cysteine residues and dephosphorylation at serine 3, Cofilin-1 loses its affinity for actin, translocates from the cytosol to mitochondria and induces apoptosis ^57,58^. Oxidation of methionine 115 apparently also prevents its actin-depolymerization activity and induces forced mitochondrial translocation and apoptosis ^59^. In addition, Cofilin-1 can be glutathionylated, which represents the presumed enzymatic activity of GDAP1. Glutathionylation of Cofilin-1 was demonstrated in lymphocytes treated with the thiol-oxidizing reagent diamide ^60^ and in cells of the rat nucleus accumbens during cued cocaine seeking in the absence of cocaine ^50^. Interestingly, glutathionylation reduces cofilin-1-dependent depolymerization of F-actin, implying regulatory functions in cell signaling ^50^. It remains to be clarified whether GDAP1 affects the redox state and function of Cofilin-1 by specific glutathionylation.

Mutations in other proteins that target the same pathway also cause CMT disease. Mutated PDK3 also inhibits the PDC by hyperphosphorylation of the PDH E1α subunit similar to GDAP1 loss of function. This causes CMTX6 ^26^. Loss of function of the actin-polymerizing protein INF2 alters mitochondrial shape by changing DRP1 access to mitochondria similar to *GDAP1* KD ^19^. INF2 is also implicated in the abundance of MERCS and changes in mitochondrial Ca^2+^ dynamics ^20^ which we also observed in GDAP1 loss-of-function cells. Mutations in INF2 also cause CMT disease, CMTDIE ^22^. Mutations in Dynamin-2 (DNM2), a ubiquitously expressed large GTPase that interacts tightly with the actin and microtubule network ^61,62^ also cause CMT disease, dominant intermediate CMT (CMTDIB) ^63^ and autosomal-dominant CMT2M ^64^. Similar to GDAP1, DNM2 works in concert with DRP1 to orchestrate mitochondrial constriction events that result in division ^65^. DNM2 therefore directly interferes with the same processes that we found to be affected by GDAP1 loss of function. Finally, mutations in BSCL2 (Bernardinelli-Seip Congenital Lipodystrophy Type 2) aka seipin cause a distal hereditary motor neuronopathy (HMN5C) which can present with features of axonal CMT2 ^66^. BSCL2 is an ER protein crucial in the formation of lipid droplets ^67,68^. BSCL2 also interacts with the adaptor protein YWHAB (14-3-3-beta) which then recruits cofilin-1 to remodel the actin cytoskeleton for adipocyte differentiation ^69^.

In summary, our results shed light on the pathophysiology of CMT4A and highlights the importance of the actin cytoskeleton for mitochondrial function. The fact that mutations in diverse proteins that all interact with the overarching mechanisms described here – metabolic and mitochondrial remodeling caused by changes in the the interaction of the actin cytoskeleton with mitochondria – cause CMT disease strengthens the importance of this mechanism for the health of the peripheral nervous system.

## Material and Methods

### Generation of iPSCs

iPSC lines from two CMT4A patients and one healthy donor were generated with Sendai virus (CytoTune, DNAVEC Corporation) coding for POU5F1, SOX2, KLF4 and MYC. All subjects gave written informed consent prior and the study was approved by the local ethics committee at the Universities of Warsaw (patient #1), Düsseldorf (patient #2) and Bonn (healthy donor). In brief, Sendai virus-infected primary fibroblasts were immediately centrifuged for 45 min at 32 °C with 1,500 g (spinfection) and cultivated in Advanced DMEM containing 5% fetal calf serum (FCS) and 1% GlutaMAX™ (all from Life Technologies). On the following day the virus-containing medium was replaced with fresh culture medium. Five days post infection (d5), transduced fibroblasts were trypsinized and seeded onto mouse feeder-coated dishes in DMEM/F-12 containing 10% KnockOut™ Serum Replacement, 1% nonessential amino acids (NEAA), 1% GlutaMAX™, 1% pyruvate, 0.1 mM β-mercaptoethanol and 10 ng/ml basic fibroblast growth factor (bFGF) (all from Life Technologies). Medium was changed every other day until clonal iPSC colonies were manually picked and adapted to feeder-free culture conditions. Several clonal lines were subjected to SNP genotyping in order to identify iPSC clones with normal karyotype.

### SNP analysis of iPSC lines

Genomic DNA was prepared using the DNeasy Blood & Tissue Kit (Qiagen). Whole-genome single nucleotide polymorphism (SNP) genotyping was performed at the Institute of Human Genetics at the University of Bonn. Genomic DNA at a concentration of 60 ng/μL was used for whole-genome amplification. Afterwards, the amplified DNA was fragmented and hybridized to sequence-specific oligomers bound to beads on an Illumina HumanOmniExpress-12v1.0 chip. Data were analyzed using Illumina Bead Studio.

### Differentiation of hiPSCs into neural cells

Induced pluripotent stem cells were grown using mTeSR™1 medium (Stemcell Tech., 05850) on matrigel (Corning, 354277). Human iPS cells were detached from matrigel using ReLeSR™ (Stemcell Tech., 05872), resuspended in medium consisting of Dulbecco’s modified Eagle’s medium/ F12 supplemented with 10% knockout serum (Invitrogen, 10828-028), 1% N2 supplement (Gibco, 17502-048), 0.05% B27 (Gibco, 17504-044), 20 ng/ml epidermal growth factor (Sigma, E9644) and 10 ng/ml basic fibroblast growth factor (Gibco, PHG0024), and plated within a low-attachment petri dish to induce embryoid body (EB) formation for 4 days. The EBs were plated on polyornithine (Sigma, P3655)/laminin (Sigma, L2020)-coated dishes for additional 6 days to induce the formation of neural rosettes. Neural rosettes were then manually removed, dissociated with accutase (Stemcell Tech., 07920), plated on poly-L-ornithine/laminin-coated dishes, and then treated with 3 μM retinoic acid for 7 days. The medium was changed daily and cultures were passaged weekly by accutase and plated on matrigel-coated dishes in the above-mentioned neural medium.

### Small molecule differentiation of hiPSCs into motoneurons

iPSCs differentiation was performed by the addition of small molecules adapted from a previously described protocol ^70^. Briefly, for neuronal induction, iPSCs were seeded as colonies resuspended in E8 medium 2 days prior differentiation. To start the differentiation, medium was changed to N2B27 medium (50% Neurobasal medium/ 50% DMEM-F12 medium with 1:200 N2 supplement, 1:100 B27 supplement without vitamin A and 1% penicillin, streptomycin and L-glutamine respectively) supplemented with the small molecules 10 μM SB-431542 (SB), 1 μM dorsomorphin (DM), 3 μM CHIR 99021 and 0.5 μM PMA for four days. Subsequently, SB and DM were replaced by 150 μM Ascorbic Acid (AA), and cells were fed daily until epithelium-like structures emerged. Neural epithelial structures were picked, dissociated mechanically and plated on 12-well plates, which were coated with Matrigel (1:100 in DMEM/F12) overnight. After around five passages the smNPC cells reached a high purity and were further kept in culture on Matrigel coated plates and N2B27 medium supplemented with CHIR, PMS and AA. Detaching of the cells for passaging was performed with accutase. Starting from smNPC passage 13, differentiation to MN could be initiated. N2B27 medium with 1 μM PMA was added 3 days after passaging. After two more days, 1 μM retinoic acid (RA) and 1 μM PMA were supplemented to the medium for the following 8 days, until culturing in maturation medium began, consisting of N2B27 medium with BDNF, GDNF and dbcAMP for two more weeks.

### Immunoblotting

Denatured total cellular protein samples were separated on SDS polyacrylamide gels 4-15% Mini-PROTEAN^®^ TGX Stain-Free™ gels (Bio-Rad) and transferred onto a nitrocellulose membrane using the Trans-Blot^®^ Turbo™ Transfer System (Bio-Rad). Membranes were blocked with 3% (w/v) milk powder in PBS-T or TBS-T (1x PBS or TBS, 0.05% (v/v) Tween 20) for 1 h at room temperature (RT). Chameleon Duo Pre-stained Protein Ladder (Li-Cor Biosciences) was used as molecular weight standard. Primary antibodies were anti-actin mAB (clone C4, 1:4000; Millipore MAB1501), anti-Cofilin-1 mAB (clone D3F9, 1:1000, Cell Signaling, 5175), anti-p-Cofilin-1 mAB (Ser3, clone 77G2, 1:1000, Cell Signaling 3313), anti-DRP1 mAB (clone 4E11B11, 1:1000, Cell Signaling 14647), anti-GDAP1 (1:750, Sigma HPA014266), anti-G6PD mAB (clone D5D2, 1:1000, Cell Signaling 12263), anti-GAPDH mAB (clone 14C10, 1:2000, Cell Signaling 2118), anti-GDH mAB (clone D9F7P, 1:1000, Cell Signaling 12793), anti-HK1 mAB (clone C35C4, 1:1000; Cell Signaling 2024), anti-HK2 mAB (clone C64G5, 1:1000, Cell Signaling 2687), anti-LDHA mAb (clone C4B5, 1:1000, Cell Signaling 3582), anti-MCU (1:1000, Sigma HPA016480), anti-MFF (1:1000, Proteintech 17090-1-AP), anti-MFN2 mAB (clone M03, 1:500; Abnova H00009927-M03), anti-Nestin mAB (clone rat-401,1:1000, Merck chemicals MAB353), anti-PDH mAb (E1α, clone C54G1, 1:1000, Cell Signaling 3205), anti-PDH mAB (E2,E3bp, clone 13G2AE2BH5, 1:1000, Abcam, ab110333) anti-p-PDH (Ser293, 1:1000, Cell Signaling 31866), anti-PKM1/2 mAb (clone C103A3, 1:1000, Cell Signaling 3582), anti-TOM20 (1:1000, Sigma HPA011562), anti-βIII-Tubulin mAB (clone TuJ-1, 1:1000, R&D Systems MAB1195). The membranes were incubated with the primary antibodies overnight at 4°C. For visualization, membranes were incubated with an infrared fluorescence IRDye^®^ 680RD Streptavidin for biotinylation staining or IRDye 800-conjugated anti-mouse, 800-conjugated anti-rabbit, or 680-conjugated anti-mouse IgG secondary antibody (1:15,000; Licor), for 1 h at RT and detected with the Odyssey Infrared Imaging System (Licor). Western Blots were analyzed with the Image Studio Lite Software (Li-Cor Biosciences).

### Immunocytochemistry (ICC)

Neuronal cells were grown on matrigel coated μ-Slide 8 Well, ibiTreat (Ibidi, 80826), fixed with 4% paraformaldehyde (PFA) (CarlROTH, 3105.2) and permeabilized by 0.25% (v/v) Triton X-100 in PBS. Unspecific binding of antibodies was blocked with 1X Roti^®^-Block (CarlROTH, A151.4) for 30 min. Primary antibodies anti-Nestin (1:100, Ebioscience, 14-9843-82), anti-MAP2 (1:500, Synaptic Systems, 188004), anti-Smi32 (Clone SMI32, 1:1000, Biolegend 801702) were treated in 1X Roti^®^-Block at 4 °C overnight. The cells were washed three times with PBS and incubated with the fluorescent conjugated secondary antibody (Molecular Probes, A-11001) in 1X Roti^®^-Block for 1 h at RT. Subsequently, three PBS washing steps were done. Nuclei were stained with 300 nM 4’,6-diamidino-2-phenylindole (DAPI). Pictures were taken with a Leica TCS SP5 inverted confocal microscope with a 63x/NA1.4 oil immersion objective, a BX51 Fluorescence microscope (Olympus) with a 20x objective or an Opera PhenixTM spinning disc high content screening microscope (Perkin Elmer, USA) equipped with two 16-bit sCMOS cameras and a 40x, 1.1 NA water immersion objective. For Cofilin-1 evaluations, SH-SY5Y cells were grown on 12 mm glass cover slides. Primary antibodies were anti-Cofilin-1 mAB (clone D3F9, 1:100, Cell Signaling, 5175), anti-GRP75 mAB (clone N52A/42, 1:20, Neuromab 75-127). Pictures were taken with a Leica TCS SP8 inverted confocal microscope with a 63x/1.4 NA oil immersion objective in z-stacks. Images were analyzed in Imaris 9.5.1 (Bitplane).

### Confocal microscopy and image analysis

High content light microscopy analysis was conducted with the Opera Phenix™ spinning disc microscope. Fluorescence (Ex/Em) for Hoechst, MitoTrackerTM Green FM and tetramethylrhodamine methyl ester perchlorate (TMRM) was measured at 405/435-480, 488/500-550 and 561/570-630 nm ex/em, respectively. BODIPY™ 558/568 C12 was added to the cells in parallel to MitoTracker in a final concentration of 1 μM for 15 min and subsequently, cells were incubated with normal growth medium. The images were analyzed with the software packages Harmony (version 4.5) and Columbus (version 2.7.1) containing the PhenoLOGICTM machine learning plugin (Perkin Elmer). The complete image analysis sequences are available upon request. Live-cell imaging of actin filament analysis was conducted two days after transfection of GFP-F-tractin (gift of Henry N. Higgs, Dartmouth University) and together with staining of mitochondria with 100 nM MitoTracker™ Red CMXRos. Images from confocal microscopy for Cofilin-1, GRP75 and actin filament quantification were analyzed in Imaris 9.5.1 (Bitplane). To improve the signal-to-noise ratio, iterative deconvolution was performed (Huygens Essential 20.10). The software then created 3D surfaces of mitochondria and Cofilin-1 spots to measure the surface area and to determine the shortest distance from the Cofilin-1 surface to the mitochondrial surface. Colocalizing Cofilin-1/actin with a distance ≤ 0 μm was duplicated to a new surface and taken for the Imaris surface-surface contact XTension to quantify the contact area, defined as the percentage of mitochondrial surface contacting Cofilin-1.

### Cell culture and generation of stable cell lines

SH-SY5Y cell lines were grown in DMEM/F12 medium (Gibco) supplemented with 10% (v/v) fetal calf serum (FCS; Thermo Scientific), 2 mM L-glutamine (Gibco), 1 × MEM non-essential amino-acids, 100 U/ml penicillin and 100 μg/ml streptomycin (Gibco) in a humidified incubator at 37°C, 5% CO2 and 95% air. The SH-SY5Y cell lines pLKO-NT (control) and G4 (knockdown) were a kind gift of David Pla-Martin and Francesc Palau ^21^ and were grown in growth medium containing 2 μg/ml puromycin (InvivoGen).

### Measurement of mitochondrial oxygen consumption

Mitochondrial respiration and oxygen consumption were analyzed using the Oxygraph-2k (Oroboros Instruments). Cells in suspension at a density of 1.5 - 2.0 × 106 cells/ml were analyzed under continuous stirring at 750 rpm and 37°C. All chemicals were purchased from Sigma. In a phosphorylation-control-protocol, the routine respiration of cells in their general growth medium was measured following the addition of 2 μg/ml oligomycin to inhibit the ATP synthase and measure leak respiration. By titration of the protonophore carbonyl cyanide 4-(trifluoromethoxy) phenylhydrazone in 0.5 μM steps the respiration was stimulated up to a maximum oxygen flow and the electron transfer system capacity was determined. By the addition of 0.5 μM rotenone and 2.5 μM antimycin A the respiration was inhibited and the non-mitochondrial residual oxygen consumption was measured. In 2 s intervals, the oxygen concentration and the oxygen flow per cells were recorded using the DatLab software 5.1 (Oroboros Instruments). All measurements were performed after daily calibration of the polarographic oxygen sensors and using instrumental background correction. The measured respiratory states were analyzed after correction with ROX to compare only mitochondrial oxygen consumption.

### ATP measurements

Cytosolic and mitochondrial ATP levels were quantified as described ^29^. Cells were transiently transfected with plasmids carrying the bioluminescence energy transfer (BRET)-based ATP biosensor BTeam without targeting signal sequence (for cyto-ATP determination) or targeted to mitochondria (for mito-ATP determination) using TurboFectin reagent (OriGene). 48 h later, the cells were incubated for 30 min in phenol red-free medium supplemented with 30 μM NanoLuciferase (NLuc) inhibitor to avoid disturbance from the BTeam released from dead cells. Afterwards, NLuc substrate (Promega) was added to the medium and the plate incubated for 20 min. Subsequently, luminescent emissions from the cells were measured at 37°C at 520/560 nm (Yellow Fluorescent Protein (YFP) emission) and at 430/470 nm (NLuc emission) using a Tecan Spark^®^ Multimode Microplate Reader.

### Glutamine and glutamate measurements

The glutamine-glutamate-glo assay (Promega, # J8021) was performed according to manufacturer’s instructions to determine intracellular glutamine and glutamate concentrations. Briefly, 20,000 cells of the SH-SY5Y cell line and 100,000 motoneurons were plated in triplicates per experiment on white 96-well plates (Greiner, # 655083) two or seven days prior to the experiment, respectively. On the day of experiment, cells were washed twice with PBS and 30 μl of PBS as well as 15 μl of 0.3 HCl solution was added to the cells and mixed for 5 min. 15 μl of 450 mM Tris solution, pH 8.0 was added and incubated for further 60 sec. From each lysate, 25 μl was transferred into a new white 96 well plate for I) glutamine plus glutamate and II) glutamate only measurement. Glutaminase was only added to the first set of wells and incubated for 30 min at RT. The detection reagent composed of Luciferin detection solution, reductase, reductase substrate, GDH and NAD was added to all wells and incubated for 60 minutes at RT before luminescence detection in a Tecan Infinite 200 pro plate reader. Concentrations were calculated with glutamine and glutamate standard curves (0.78 μm-50 μM) and a blank was included to remove any assay background signal. Glutamate levels were calculated by subtracting the glutamate-only signal from the signal containing glutamate and glutamine levels together.

### Electron microscopy and analysis

Cells were pelleted by centrifugation at 1300 rpm for 3 min and fixed in 3% glutaraldehyde overnight. Following several rinses in 0.2 M sodium cacodylate buffer (pH 7.3), the samples were postfixed in 1% osmium tetroxide in cacodylate buffer for 2 h, dehydrated through an ascending series of ethanol concentrations and embedded in resin with propylene oxide as an intermediary. Semi-thin (0.65 μm) sections for light microscopy and ultrathin (50 nm) sections for electron microscopy were cut on a Leica Ultracut UCT ultramicrotome. Semi-thin sections were stained with methylene blue. Ultrathin sections were stained with an alcoholic solution of 1% uranyl acetate and lead citrate in sodium hydroxide and examined with a Zeiss EM-910 transmission electron microscope. For morphometric and quantitative analysis, representative cells were photographed at a magnification of 10,000 and 18,000. Analyses were done with ImageJ. MERCs length and width were analyzed considering the contact site in a range of 0-110 nm. The frequency distribution of the MERCs width was calculated using GraphPad.

### Biochemical analysis of mitochondria-ER contact sites

Cells were detached using trypsin/ETDA and resuspended in 1-2 ml of a buffer containing 0.32 M sucrose, 10 mM Tris-HCl, 1 mM EDTA and protease (Roche Diagnostics, 04693124001) and phosphatase (Roche Diagnostics, 04906845001) inhibitors. Cells were disrupted by a nitrogen decompression instrument (Parr Instrument Company, 4639) and centrifuged (2,000 g, 10 min). The supernatant was transferred to a new microtube, centrifuged at 10,000 g for 10 min and the cell pellet re-suspended in 1 ml of the same buffer containing the same components as above except 0.5 M sucrose. Finally, mitochondria were sedimented by centrifugation at 10,000 g for 10 min and resolved in an appropriate amount of RIPA buffer for the subsequent experiments.

### FRET-based FEMP probe to quantify mitochondria-ER contact sites

MERCS were quantified with a FRET-based sensor indicating the proximity between the ER and mitochondria. The plasmid encodes for a YFP-linked outer mitochondrial membrane protein Akap1 and a CFP-conjugated ER-protein Sac1, as well as a fused FKBP and FRB domain respectively. These domains can form heterodimers upon rapamycin treatment. The specific localization of these proteins is ensured by the introduction of a self-cleavable Tav2A sequence ^36,37^. The FEMP plasmid (FRET-based ER-mitochondria probe) was transfected with GenJet. 48 h later, images were taken with the Perkin Elmer Operetta High-Content Imaging System acquiring the CFP-(410-430/460-500 ex/em), YFP-(490-510/520-560 ex/em) and YFPFRET-emission (410-430/520-560 ex/em) with a 40x water objective for determination of the basal distances between the organelles. Subsequently, the cells were treated with 100 nM rapamycine for 15 min for FKBP-FRB dimerization induction and to reach a maximum of YFP_FRET_ signal. Cells were fixed for another 20 min with 1% PFA and imaging was performed again with equal microscopy settings. For the analysis, the Harmony software was used. First, the cells were identified using the YFP channel. Within each cell and region of interest (ROI), the intensities of the three acquired channels were calculated, including background subtraction. FRET basal and FRET max were calculated as: (FYFP-FRET_cell_-FYFP-FRET_background_)/ (FCFP_cell_-FCFP_background_); FRET Ratio was calculated as (FRET_max_–FRET_basal_)/FRET_basal_.

### Mitochondrial Ca^2+^ measurement

Dye: Cells were plated (20,000 cells/cm^2^) in 8-well μ-slides ibiTreat (ibidi, 80826) and treated with 5 μM of Rhod2-AM (Molecular Probes, R1245MP) in culture medium without FBS for 60 min at 37°C the next day. Rhod2-AM fluorescent signals were analyzed at (Ex/Em) 549/578 nm wavelengths using a Leica SP5 confocal microscope and analyzed by ImageJ. Genetically-encoded reporter: Cells were transfected with the genetically-encoded reporters and analyzed two days later in an inverted TCS-SP5 confocal microscope (Leica) with appropriate excitation/emission wavelengths as reported for CEPIA3mt (Addgene, 58219) and GEM-CEPIA1er (Addgene, 58217) ^33^. The CEPIA3mt construct was co-transfected and normalized to mito-TurboFarRed.

### Aequorin Ca^2+^ measurements

For Ca^2+^ measurements, the biosensor aequorin (AEQ) was used, which is a 22 kDa calcium-binding photoprotein isolated from jellyfish Aequorea Victoria ^71^. SH-SY5Y cells were grown on 12 mm glass coverslips to a confluence of 40-50% and transfected with cytosolic or mitochondria-targeted AEQ (cytAEQ/mtAEQ) using GenJet. On the day of the experiment, the cells were treated with 5 μM coelenterazine-N-AM (Santa Cruz Biotechnology sc-205904) in basic saline solution (BS, 135 mM NaCl2, 5 mM KCl, 0.4 mM KH2PO4, 1mM MgSO4 × 7 H2O, 20 mM HEPES, 0.1% (w/v) glucose, pH 7.4 adjustment at 37°C with NaOH 10 N) containing 1 mM CaCl2 for 2 h at 37°C and 5% CO2. Coelenterazine served as a substrate for AEQ. Cover slides were then placed in the luminometer with constant buffer perfusion with BS supplemented with I) 1 mM Ca^2+^ (30 s) II) 200 μM EGTA (30 s) III) 200 μM Carbachol (CCH) + 200 μM EGTA (120 s) IV) 100 μM Digitonin + 5 mM CaCl2 (220 s).

### Precipitation of biotinylated Avi-GDAP1

Primary cortical neuron cultures were prepared from embryos (E16) from the transgenic mouse line Gt(ROSA)26Sortm1(birA)Mejr (ROSA26-BirA) of a C57BL/6N background. In this mouse strain, the biotin ligase BirA was inserted into the gene locus of the ROSA26 promotor ^41^. Cortical neurons were cultured on poly-D-lysine coated plates (0.05 mg/ml) in Neurobasal medium (NBM, Life Technologies, 21103049) supplemented with 2% (v/v) B-27 supplement (Life Technologies, 17504-044), 1% (v/v) L-glutamine (Sigma-Aldrich, G1251) and 100 U/ml penicillin and 100 μg/ml streptomycin (Sigma-Aldrich, P0781). Medium change was performed at day 1 and day 4 after isolation. For transduction of the primary neurons, viruses were added in a volume that equaled 6*10^7^-8*10^7^ copies/μl per well of a 6 well plate containing 4 ml of NBM and neurons on day 4 after isolation. Neurons were cultured for further 7 days. Cells were harvested, washed with PBS and lysed in RIPA buffer supplemented with protease inhibitors.

The samples were centrifuged at 21,000 g for 30 min at 4°C, the proteins concentration was determined via BC-Assay and 20 μg protein lysate was removed as input-control. Protein lysates were either directly incubated with Dynabeads™ MyOne™ Streptavidin T1 according to the manufacturer’s protocol or taken for GSH or GSSG incubation (5 mM GSH or GSSG in 10 mM HEPES pH 7.4, 35 mM sucrose, 40 mM KCl, 0.25 mM EGTA, 2 mM Mg(CH3COO)2, 0.5 mM GTP, 1 mM ATP (K^+^), 5 mM Na-succinate, 0.08 mM ADP, 2 mM K2HPO4 for 30 min at 37°C). Dynabeads™ MyOne™ Streptavidin T1 were incubated for 45 min at RT on a shaker. Dynabeads were washed three times with RIPA buffer and once with PBS and stored at −80°C until further processing for LC-MS.

### Protein digestion

Bound proteins were eluted from Dynabeads in 10 mM Tris pH 8.0, 2% SDS, 1 mM Biotin at 80°C and digested according to the SP3 (“Single-Pot Solid-Phase-Enhanced Sample Preparation’’) protocol ^72^. Proteins were reduced by adding 5 μl of 200 mM Dithiothreitol (DTT) per 100 μl lysate (45°C, 30 min). Free cysteines were subsequently alkylated by adding 10 μl 100 mM Iodoacetamide (IAA) per 100 μl lysate (RT, 30 min, in the dark). Subsequently, remaining IAA was quenched by adding 10 μl 200 mM DTT per 100 μl lysate. Magnetic carboxylate modified particles Beads (SpeedBeads, Sigma) were used for protein clean-up and acetonitrile (ACN), in a final concentration of 70%, was added to the samples to induce the binding of the proteins to the beads by hydrophilic interactions (18 min RT). By incubating the bead-protein mixture on a magnetic stand for 2 min, the sample was bound to the magnet and the supernatant removed, followed by two washing-steps with 70% ethanol (EtOH), addition of 180 μl ACN, incubation for 15 s and removal of the solvent. Finally, 5 μl digest buffer (50 mM ammonium bicarbonate, 1:25 w/w trypsin:protein ratio) was added to the air-dried bead-protein mixtures and incubated overnight at 37°C. To purify peptides after digestion, ACN was added to a final concentration of 95%. After another washing step. the beads were resuspended in 10 μl 2% DMSO (in water), put into an ultrasonic bath for 1 min and then shortly centrifuged. 10 μl of the resulting supernatant was mixed with 5 μl 100 fmol/μl Enolase digest (Waters Corporation) and acidified with 5 μl 1% formic acid (FA).

### LC-MS analysis

Liquid chromatography (LC) of tryptic peptides was performed on a NanoAQUITY UPLC system (Waters Corporation) equipped with 75 μM × 250 mm HSS-T3 C18 column (Waters corporation). Mobile phase A was 0.1% (v/v) formic acid (FA) and 3% (v/v) DMSO in water. Mobile phase B was 0.1% (v/v) FA and 3% (v/v) DMSO in ACN. Peptides were separated running a gradient from 5 to 40% (v/v) mobile phase B at a flow rate of 300 nL/ min over 90 min. The column was heated to 55°C. MS analysis of eluting peptides was performed by ion-mobility enhanced data-independent acquisition (UDMS^E^) ^40^. Precursor ion information was collected in low-energy MS mode at a constant collision energy of 4 eV. Fragment ion information was obtained in the elevated energy scan applying drift-time specific collision energies. The spectral acquisition time in each mode was 0.7 s with a 0.05 s interscan delay resulting in an overall cycle time of 1.5 s for the acquisition of one cycle of low and elevated energy data. [Glu1]-fibrinopeptide was used as lock mass at 100 fmol/μL and sampled every 30 s into the mass spectrometer via the reference sprayer of the NanoLockSpray source. All samples were analyzed in three technical replicates.

### Data processing and label-free protein quantification

UDMS^E^ data processing and database search was performed using ProteinLynx Global Server (PLGS, ver. 3.0.2, Waters Corporation). The resulting proteins were searched against the UniProt proteome database (species: *Mus musculus,* UniProtK-Swissprot release 2019_05, 17.051 entries) supplemented with a list of common contaminants. The database search was specified by trypsin as enzyme for digestion and peptides with up to two missed cleavages were included. Carbamidomethyl cysteine was set as fixed modification and oxidized methionine as variable modification. False discovery rate assessment for peptide and protein identification was done using the target-decoy strategy by searching a reverse database and was set to 0.01 for database search in PLGS. Retention time alignment, exact mass retention time (EMRT), as well as normalization and filtering were performed in ISOQuant ver.1.8. ^40,73^. By using TOP3 quantification, absolute in-sample amounts of proteins were calculated.

### Statistical analysis

Outliers were removed by the ROUT test and normal distribution tested using the Shapiro-Wilk test. Statistical significance was then verified using appropriate parametric (Student *t*-test or ANOVA) or non-parametric tests (Mann Whitney and Kruskal Wallis tests) followed by multiple comparison tests. The Wilcoxon signed rank test was used when normalization to 100% was necessary as indicated. Statistical analysis of mass spectrometry data was performed using two-tailed, paired *t*-tests and subsequent Bonferroni correction. Here, a corrected *p*<0.01 was considered significant, in all other data a *p*<0.05 was considered to be statistically significant.

## Acknowledgment

The excellent support by the IMB Core Facility Microscopy is gratefully acknowledged. This work was supported by grants of the Deutsche Forschungsgemeinschaft to AM (CRC1080 project A10) and to AM and EMH and ST (DFG ME1922 16-1; Ha 8334/2-2; TE599/6-1) within the priority program 1710. This work was supported by an Excellence Grant of the Luxembourg National Research Fund (FNR) within the PEARL programme (FNR; FNR/P13/6682797 to RK). Work of RK is further supported by a grant from the FNR within the National Centre of Excellence in Research on Parkinson’s disease (NCER-PD). We thank the Disease Modelling and Screening Platform of the Luxembourg Institute of Systems Biomedicine and the Luxembourg Institute of Health for their support on high content imaging of motoneurons. We thank Marion Silies for excellent and exhaustive proofreading.

## Author contribution

CW, AP, SB, AnnP, DB, FdB, CV, IB, carried out experiments; KD, MP and OB generated the iPS cells; JS and SR helped establishing high-content microscopy; SA and ST analyzed quantitative proteomic analyses; MG supervised and analyzed the FEMP and aequorin measurements; RK supervised and analyzed the experiments with human motoneurons; CW and EMH analyzed data and helped writing the manuscript; AM devised experiments, analyzed data and wrote the manuscript.

## Conflict of Interest Statement

The authors declare that they have no conflict of interest.

## Abbreviations

CMT: Charcot-Marie-Tooth
DRP1: Dynamin-related protein 1
GDAP1: ganglioside-induced differentiation associated protein 1
AR: axonal-recessive
ER: endoplasmic reticulum
MERCS: mitochondria-ER contact sites
GST: glutathione-transferase
iPSCs: induced pluripotent stem cells
NPCs: neural precursor cells
Δψm: mitochondrial membrane potential
TMRM: tetramethylrhodamine methyl ester
KD: knockdown
ETS: electron transfer system
netR: phosphorylating respiration
TEM: transmission electron microscopy
bFGF: basic fibroblast growth factor
FCS: fetal calf serum
RT: room temperature
ICC: Immunocytochemistry
PBS-T: PBS-Tween 20
Ex: Excitation
Em: Emission
PCP: phosphorylation-control-protocol
R: routine respiration
Omy: oligomycin
L: leak respiration
FCCP: carbonyl cyanide 4-(trifluoromethoxy) phenylhydrazone
E: ETS capacity
Rot: rotenone
Ama: antimycin A
ROX: non-mitochondrial residual oxygen consumption
BTeam: BRET-based ATP biosensor
NLuc: NanoLuciferase
2DG: 2-Deoxy-D-Glucose
YFP: Yellow Fluorescent Protein
CRISPR-Cas9: Clustered Regularly Interspaced Short Palindromic Repeats - CRISPR-associated protein 9
gRNA: guided-RNA
NeoR: Neomycin resistance gene
SDS: sodium dodecyl sulfate
SDS-PAGE: sodium dodecyl sulfate polyacrylamide gel electrophoresis
DTT: Dithiothreitol
SD: standard deviation
SEM: standard error of the mean
Ctrl: control
OXPHOS: oxidative phosphorylation
LDHA: lactate dehydrogenase A
GDH: glutamate dehydrogenase
TCA: tricarboxylic acid cycle
PDH: Pyruvate dehydrogenase
PDC: Pyruvate dehydrogenase complex
PDK3: pyruvate dehydrogenase kinase isoenzyme 3
MCU: Mitochondrial Calcium uniporter
FRET: Förster resonance emission transfer
IP3R: inositol trisphosphate receptors

## Supplementary figure

**Supplementary Figure 1.**
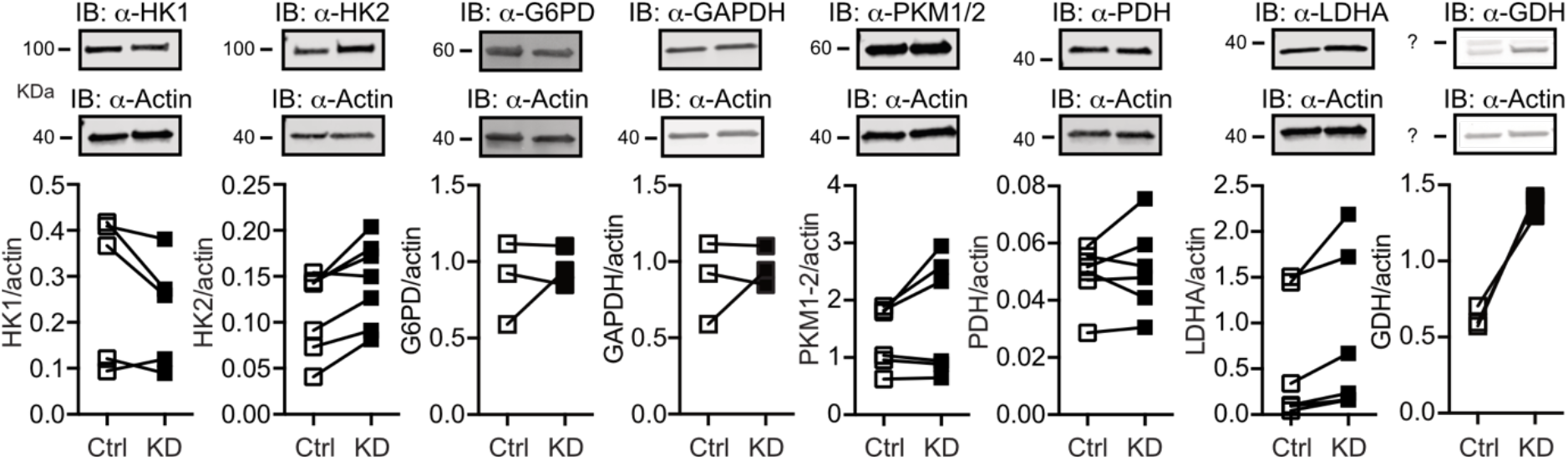
Changes in protein abundance. **(a)** Immunoblot of whole cell lysates from Ctrl and GDAP1 KD cells against Hexokinase 1 (HK1), HK2, Glucose-6-phosphate dehydrogenase (G6PD), Glyceraldehyde 3-phosphate dehydrogenase (GAPDH), Pyruvate kinase M1/2 (PKM1/2), Pyruvate dehydrogenase (PDH), Lactate Dehydrogenase A (LDHA), Glutamate dehydrogenase (GDH). Sizes are indicated and actin served as loading control. n=3-7

## Notes

### Competing Interest Statement

The authors have declared no competing interest.

